# SARS-CoV-2 ORF7a mutation found in BF.5 and BF.7 sublineages impacts its functions

**DOI:** 10.1101/2023.09.06.556547

**Authors:** Uddhav Timilsina, Sean R. Duffy, Arnon Plianchaisuk, The Genotype to Phenotype Japan (G2P-Japan) Consortium, Jumpei Ito, Kei Sato, Spyridon Stavrou

## Abstract

A feature of the SARS-CoV-2 Omicron subvariants BF.5 and BF.7 that recently circulated mainly in China and Japan was the high prevalence of ORF7a: H47Y mutation. Here we evaluated the effect of this mutation on the three main functions ascribed to SARS-CoV-2 ORF7a protein. Our findings show that H47Y mutation impairs the ability of SARS-CoV-2 ORF7a to antagonize type-I interferon (IFN-I) response and to downregulate Major Histocompatibility Complex-I (MHC-I) cell surface levels, but had no effect in its anti-SERINC5 function. Overall, our results suggest that the H47Y mutation of ORF7a affects important functions of this protein resulting in changes in virus pathogenesis.

**Importance:** In late 2021, the Omicron (B.1.1.529) VOC emerged and outcompeted the circulating VOC Delta (B.1.617.2). Soon afterwards, Omicron VOC has expanded and diversified in the world. Among the emerged subvariants of Omicron are BF.5 and BF.7 that are characterized by the presence of a number of mutations across their genome and spread quite effectively across China and other countries later on. One such mutation that was found in the vast majority of isolates of the BF.5 and BF.7 subvariants was ORF7a: H47Y, whose effect on ORF7a is unknown. In this report, we show that H47Y inhibits a number of ORF7a functions, which can potentially affect virus pathogenesis.

## INTRODUCTION

Severe acute respiratory syndrome coronavirus 2 (SARS-CoV-2) has led to a global pandemic of coronavirus disease 2019 (COVID-19). SARS-CoV-2 genome encodes four structural proteins (Spike, S; Membrane, M; Envelope, E; and Nucleocapsid, N) and a number of non-structural proteins involved in virus replication (nsp1 to nsp16) or accessory proteins involved in the modulation of host responses (ORF3a, ORF3b, ORF6, ORF7a, ORF7b, ORF8, ORF9b, ORF9c, and ORF10) (1, 2). Over the course of the COVID-19 pandemic, the continued evolution of SARS-CoV-2 has led to the emergence of several variants (Alpha, Beta, Gamma, Delta, Epsilon, Eta, Ota, Kappa, Omicron, Zeta and Mu) (3–5). The rise of SARS-CoV-2 variants of concern (VOCs) are characterized primarily by the emergence of mutations within the S protein. S mutations have led to altered virus biology facilitating evasion of vaccine- and infection-induced immunity leading to enhanced transmissibility of SARS-CoV-2 VOCs (5–7). In addition, emerging variants have mutations across other structural proteins (E, M, N), nonstructural proteins (NSP1, NSP3, NSP4, NSP5, NSP6, NSP12) and accessory proteins ORF3a, ORF6, ORF7a, ORF8 or ORF10 (8, 9). It has been reported that new SARS-CoV-2 subvariants of Omicron, BF.7 (BA.5.2.1.7) initially prevalent in China and BF.5 dominant in Japan at the late 2022 have a unique non-synonymous mutation (H47Y) in ORF7a protein (10, 11).

SARS-CoV-2 ORF7a is a type-1 transmembrane protein with 121 amino acid residues, consisting of an N-terminal signaling region (residues 1-15), an immunoglobulin-like (Ig-like) ectodomain consisting of seven β-strands (strands A to G; residues 16-96), a transmembrane domain (TM) (97-116) and a C-terminal ER-retention motif (residues 117-121) (12). Major functions ascribed to SARS-CoV-2 ORF7a during infection include impairing the antiviral effect of host factors including Serine Incorporator 5 (SERINC5) (13) and BST2/tetherin (14), inhibiting the type I interferon (IFN-I) response (15, 16) and downregulating the levels of Major Histocompatibility Complex-I (MHC-I) (17, 18) on the cell surface. In addition, SARS-CoV-2 has been associated with the induction of autophagy (19) and apoptosis (20), and upregulation of inflammatory responses (12, 21).

SARS-CoV-2 ORF7a acts as a viral antagonist of SERINC5 by preventing its incorporation into nascent virions (13). In addition, SARS CoV-2 ORF7a interacts with S protein and SERINC5 thereby counteracting SERINC5-mediated restriction of SARS-CoV-2 infectivity (13). In addition, SARS-CoV-2 ORF7a is implicated in the evasion of the host immune response by antagonizing the type I interferon (IFN) response. SARS-CoV-2 ORF7a hijacks the host ubiquitin system to polyubiquitinate itself at K119 amino acid residue thereby blocking the IFN-α-mediated phosphorylation of Signal Transducer and Activator of Transcription 2 (STAT2) (15, 16). Furthermore, recent studies have determined that ORF7a interferes with the antigen presentation ability of host cells by interacting with the heavy chain of MHC-I, thereby disrupting the assembly of MHC-I peptide loading complex (PLC) in the endoplasmic reticulum (ER) and preventing export of peptide loaded MHC-I complexes to the cell surface (17, 18).

In this study, we elucidate the impact of H47Y mutation found in the BF.5 and BF.7 sublineages on SARS-CoV-2 ORF7a functions. We show that SARS-CoV-2 ORF7a: H47Y mutation does not affect its ability to counteract the antiviral effect of SERINC5 but inhibits the ability of SARS-CoV-2 ORF7a to antagonize IFN-I response and down-regulate MHC-I cell surface levels.

## RESULTS

### SARS-CoV-2 ORF7a: H47Y mutation

Global genomic surveillance during the COVID-19 pandemic has shown that deletion and substitution mutations in the SARS-CoV-2 *ORF7a* gene are frequent (22–25). Though no study has determined the impact of these mutations in the clinical context, certain SARS-CoV-2 *ORF7a* deletion mutations are known to impair virus replication *in vitro* (22, 26). A SARS-CoV-2 strain with a truncated *ORF7a* (115 nucleotide deletion) is found to be defective in suppressing the host immune response (22). A mutation (A105V) in the TM domain of SARS-CoV-2 ORF7a that improved its stability was associated with severe disease outcome among a group of Romanian COVID-19 patients (24). Moreover, we and others have shown that *in vitro* deletion of *ORF7a* gene reduces replication of synthetic recombinant SARS-CoV-2 virus in a cell-type specific manner (13, 27), suggesting an important role of this protein in virus replication and pathogenesis.

Since late 2021 when the first Omicron variant (B.1.1.529) was reported, multiple Omicron sub-variants (BA.1, BA.2, BA.3, BA.4, BA.5) that have emerged were classified as variants of concern (VOC) till recently (4, 28). Omicron variants possess unique S mutations within the receptor binding domain (RBD) which are key for the evasion of neutralizing antibodies (6, 9, 29, 30). Omicron BF.7 (BA.5.2.1.7) subvariant has been circulating in numerous countries since mid-2022 (11). High prevalence of a sublineage of BF.7 (named BF.7.14 by PANGO) with three additional unique mutations (ORF7a: H47Y, NSP2: V94L, and S: C1243F) were reported in China (10). Nevertheless, ORF7a: H47Y mutation had been detected in isolates from Oceania and North America early in 2020 and in the Omicron BF.5 lineage in Japan at the end of 2022 (11, 31). To trace the occurrence of ORF7a: H47Y substitution along the evolutionary history of the BA.5 lineage, we reconstructed the phylogenetic tree of 5,226 SARS-CoV-2 isolates in the BA.5 lineage (Fig. 1A) using at most ten randomly selected genomic sequences per each of 523 BA.5 subvariants and performed the reconstruction of the ancestral state of ORF7a: H47Y substitution. We found two ORF7a: H47Y substitution occurrence events in the BA.5 lineage. The first one occurred in the most recent common ancestor (MRCA) of BF.5 subvariants, and the second one occurred in the MRCA of BA.7.14 and BF.7.27 subvariants (Fig. 1A). In addition, a different mutation in the 47^th^ residue of ORF7a (H47N) was reported in South Korea early in 2020(32). It is noteworthy that H47Y mutation was also reported in SARS-CoV ORF7a (33). Nevertheless, the H residue at the 47^th^ amino acid position of ORF7a is conserved in all genomic sequences of 78 sarbecoviruses we analyzed, including those of SARS-CoV-1, SARS-CoV-1-related, SARS-CoV-2, and SARS-CoV-2-related viruses (34) (Fig. 1B). Sporadic substitutions at this position among SARS-CoV/SARS-CoV-2 viruses indicate that these changes may affect protein functions.

**Fig. 1.**
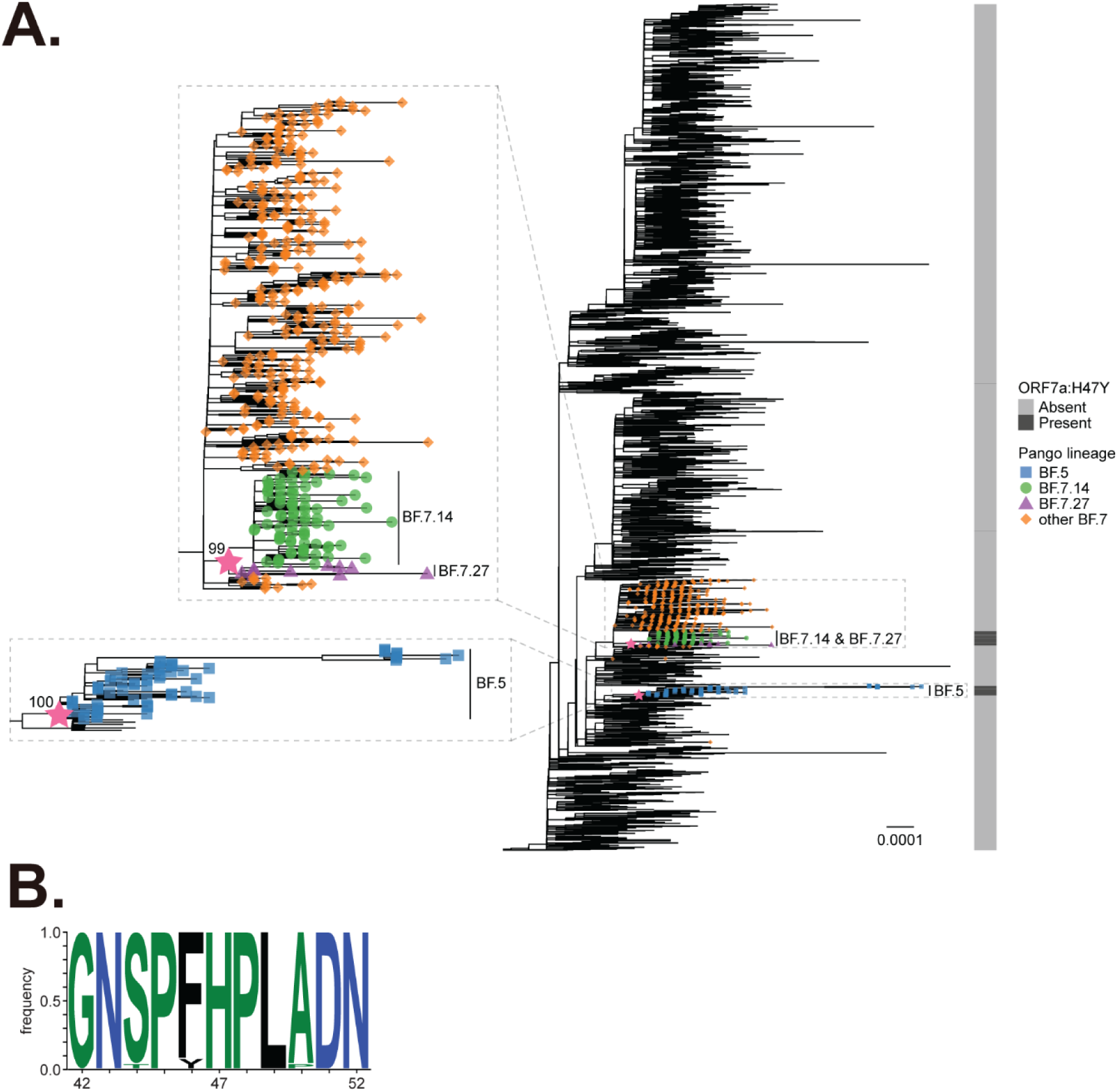
SARS-CoV-2 ORF7a H47Y mutation is prevalent among Omicron sub-lineages BF.5 and BF.7 and is highly conserved among sarbecoviruses. **(A)** A phylogenetic tree of SARS-CoV-2 in the BA.5 lineage. At most ten genomic sequences per BA.5 subvariants were randomly selected for phylogenetic tree reconstruction resulting in 5,226 sequences in total. Only BF.5 and BF.7 subvariants are labeled. The ultrafast bootstrap values of the MRCA of BF.5 and that of BF.7.14 and BF.7.27 are 100 and 99 respectively. A star represents the occurrence of ORF7a: H47Y substitution. Only the occurrence of ORF7a: H47Y substitution at an internal node with at least 5 descendant tips harboring the ORF7a:H47Y substitution is shown. **(B)** A sequence logo plot showing an amino acid frequency in ORF7a of 78 sarbecoviruses from position 42 to 52.

### SARS-CoV-2 ORF7a: H47Y mutation has no effect on its ability to counteract SERINC5

It was previously shown that SERINC5 becomes incorporated in nascent SARS-CoV-2 virions resulting in virus entry inhibition by interfering with SARS-CoV-2 S-mediated fusion (13). We also found that SARS-CoV-2 ORF7a alleviated SERINC5-mediated restriction of viral infectivity by preventing SERINC5 incorporation in nascent virions as well as forming a complex with S and SERINC5 (13). To examine the effect of the H47Y mutation of ORF7a in its ability to counteract the antiviral effect of SERINC5 on SARS-CoV-2 entry, we generated SERINC5 containing SARS-CoV-2 S containing pseudotyped viruses in the presence of either wildtype (WT) or H47Y mutant ORF7a by co-transfecting HEK 293T cells using a replication-defective HIV-1 proviral luciferase reporter plasmid (pHIV-1_NL_ΔEnv-NanoLuc), SARS-CoV-2 S. SERINC5 and either WT or H47Y ORF7a. Initially, we verified that H47Y mutation had no effect on the steady state expression level of ORF7a protein (Fig. 2A). We then utilized these pseudoviruses to infect HEK 293T-hACE2 cells and luciferase levels were measured 48 h post infection (hpi). As expected, the presence of SERINC5 reduced the infectivity of SARS-CoV-2 S pseudoviruses (Fig. 2B). In the case of the ORF7a H47Y mutant, we observed that the mutant ORF7a counteracted the SERINC5 antiviral effect on virion infectivity similar to WT ORF7a (Fig. 2B). Therefore, we conclude that the H to Y change at amino acid 47 of ORF7a does not interfere with the ability of ORF7a to counteract the SERINC5 antiviral function.

**Fig. 2.**
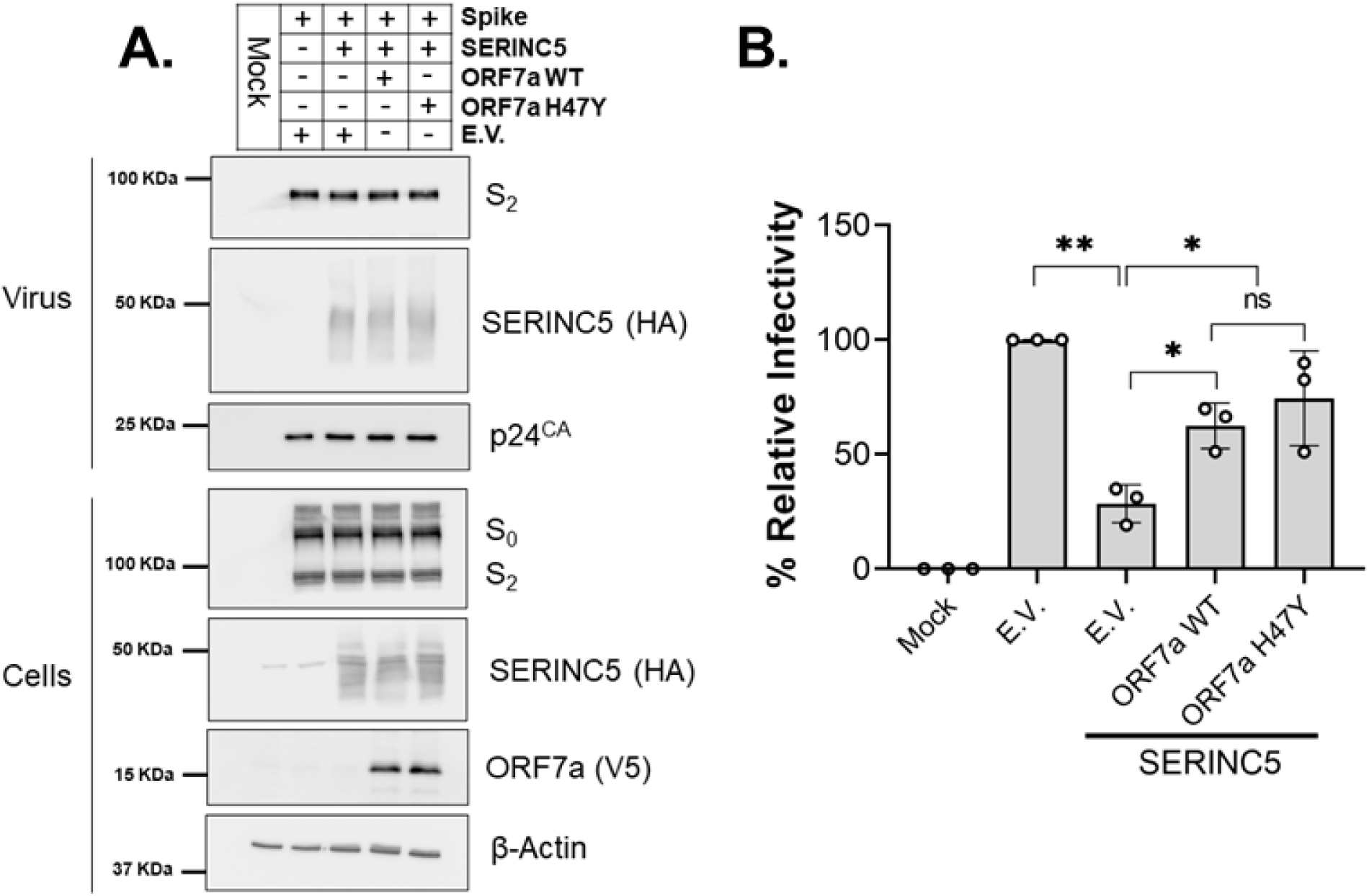
SARS-CoV-2 ORF7a: H47Y mutant retains the anti-SERINC5 activity. **(A)** SARS-CoV-2 ORF7a: H47Y mutation does not affect its steady-state expression or SERINC5 incorporation in viral particles. HEK 293T cells were co-transfected with plasmids for HIV-1_NL_ΔEnv-NanoLuc, SARS-CoV-2 Spike, SERINC5, SARS-CoV-2 ORF7a wildtype (WT)/H47Y or empty vector (E.V.) as indicated. Forty-eight hours post transfection, the indicated proteins were analyzed by immunoblotting in cell lysates and culture supernatants. Representative immunoblot images from n=3 independent experiments are shown. (**B**) SARS-CoV-2 ORF7a: H47Y mutant counteracts the antiviral effect of SERINC5. HEK 293T-hACE2 cells were infected with SARS-CoV-2 S pseudovirus from (**A**) and luciferase levels were measured 48 hpi. The percentage (%) of relative infectivity with respect to pseudovirus produced in the presence of E.V. is shown. Results are presented as mean ± SD from n=3 independent experiments. Statistical comparisons were performed by one-sample t-test (two-tailed) between E.V. and SERINC5 + E.V. conditions and unpaired t-test (two-tailed) between SERINC5 + E.V. and SERINC5 + ORF7a WT/H47Y conditions. *, P < 0.05; **, P < 0.01; ns, not significant.

### SARS-CoV-2 ORF7a: H47Y is unable to block the type I IFN response

Previous studies have reported that SARS-CoV-2 ORF7a inhibits type I IFN signaling by targeting STAT2 phosphorylation (15, 16). We utilized an ISG56-promoter-driven luciferase assay to compare the ability of ORF7a WT and H47Y mutant to inhibit the type I IFN response. HEK 293T cells expressing ORF7a WT or H47Y mutant proteins were treated with IFN-β followed by measurement of luciferase levels as a marker of ISG56 promoter activity. In agreement with previous reports, we found that the presence of WT ORF7a significantly inhibited the IFN-β-mediated activation of the ISG56 promoter (Fig. 3). However, ORF7a H47Y mutant has no effect on the ISG56 promoter activity in the presence of IFN-β, similar to what is observed in the presence of empty vector (E.V.) (Fig. 3). In summary, we show that the H47Y mutation of ORF7a interferes with the ability of ORF7a to inhibit type I IFN signaling.

**Fig. 3.**
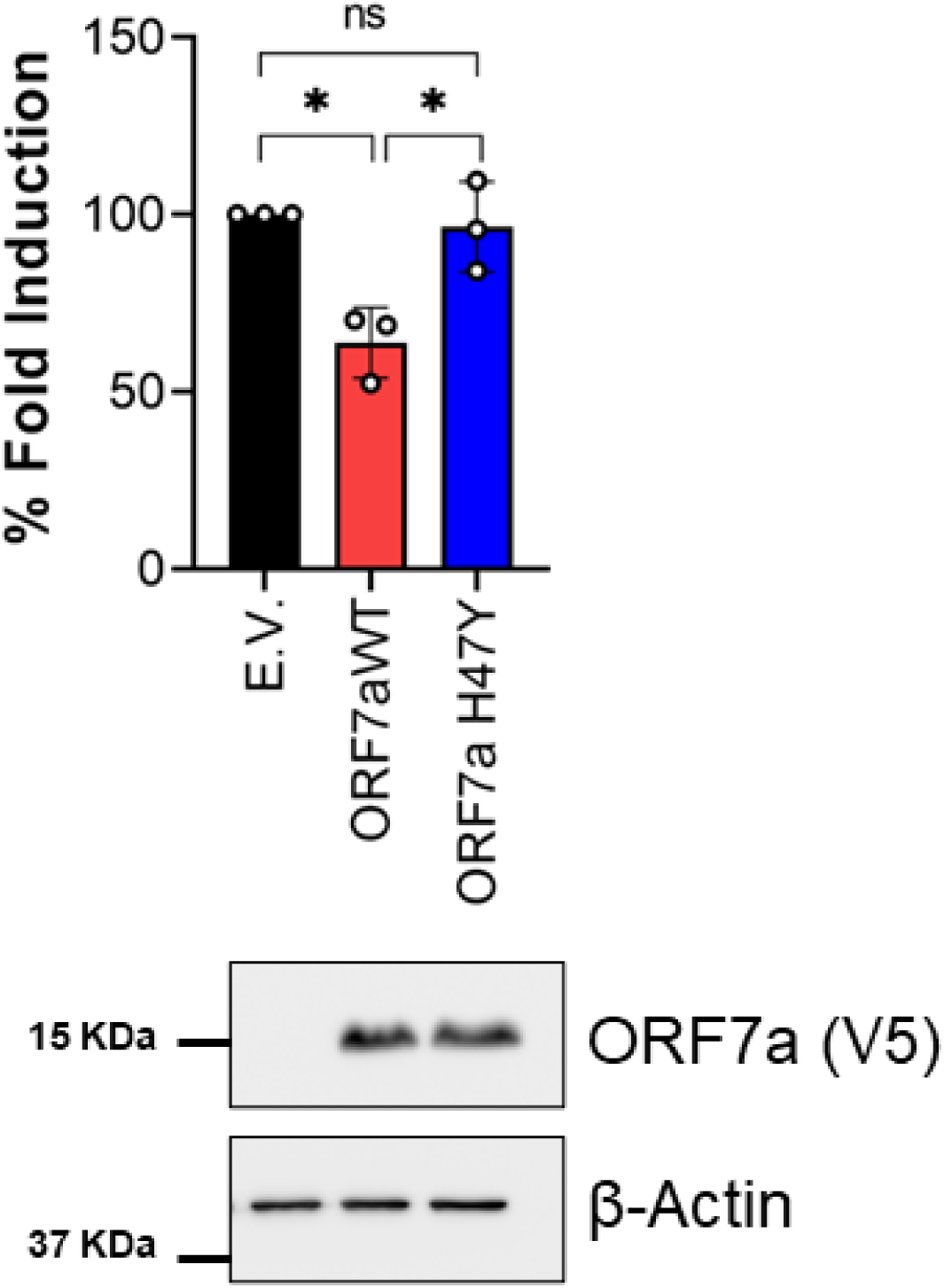
SARS-CoV-2 ORF7a: H47Y mutant is ineffective in antagonizing IFN-I response. HEK 293T cells were co-transfected with plasmids for an ISG56-promoter driven Firefly luciferase reporter, Renilla luciferase control, and SARS-CoV-2 wildtype (WT)/H47Y or empty vector (E.V.). Sixteen hours post transfection, the cells were either treated with or without human IFN-β. After eight hours of treatment, cells were assayed for dual-luciferase activities. Data were analyzed by normalizing Firefly luciferase to Renilla luciferase activities and then normalizing to non-IFN-β-treated samples to obtain fold induction. The value of the E.V. was set to 100% fold induction. Error bars represent mean ± S.D for n=3 independent experiments. One sample t-test (two-tailed) was used for comparisons between E.V. and ORF7: WT/H47Y conditions, while unpaired t-test (two-tailed) was used for comparisons between ORF7a WT and H47Y conditions. Shown below are representative immunoblot images (n=3) for verifying ORF7a protein expression in transfected cells. Cell lysates from mock-treated conditions were harvested at 24 h post transfection. *, P < 0.05; ns, not significant.

### SARS-CoV-2 ORF7a: H47Y mutation abrogates its potential to down-regulate surface MHC-I

Another major function of SARS-CoV-2 ORF7a is to physically interact and retain MHC-I complexes in the ER preventing their transport to the cell surface (17, 18). Thus, we examined the impact of H to Y substitution at position 47 of ORF7a in preventing the transport of MHC-I to the cell surface. We transfected HEK 293T cells with either ORF7a WT, ORF7a H47Y or E.V. and measured MHC-I surface levels 24 h post transfection by flow cytometry. We observed that surface levels of MHC-I were reduced in cells expressing WT ORF7a when compared to those transfected with E.V. (Fig. 4B). Interestingly, unlike WT ORF7a, ORF7a H47Y mutant did not alter the surface MHC-I levels (Fig. 4B). Thus, we conclude that H47Y mutation in SARS-CoV-2 ORF7a protein disrupts its ability to downregulate MHC-I surface levels.

**Fig. 4.**
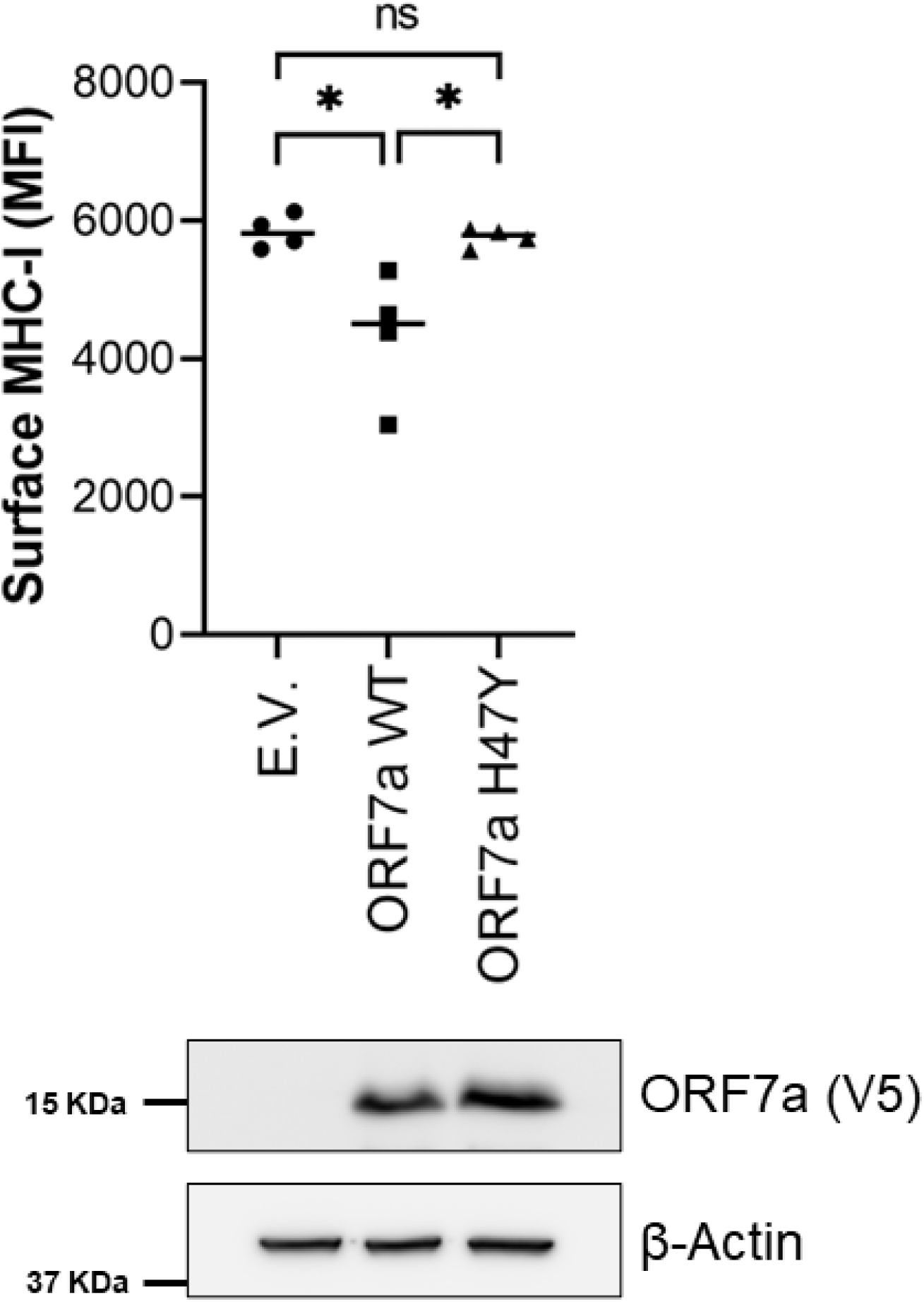
SARS-CoV-2 ORF7a: H47Y mutation impairs its ability to down-regulate cell surface MHC-I levels. HEK 293T cells were co-transfected with plasmids for eGPF and ORF7a wildtype (WT)/H47Y or empty vector (E.V.). At 24 h post transfection, cells were stained with a pan-HLA-ABC antibody (W6/32) conjugated with Alexa Flour 647 followed by flow cytometry. GPF positive cells were gated and compared for MHC-I surface levels (median fluorescent intensity). Error bars represent mean ± S.D for n=4 independent experiments. Statistical comparisons were performed using unpaired t-test (two-tailed). Shown below are representative immunoblot images (n=4) for verifying ORF7a protein expression in transfected cells. Cell lysates in transfected cells were harvested at 24 h post transfection. *, P < 0.05; ns, not significant.

### The H47Y mutation in SARS-CoV-2 ORF7a causes altered MHC-I interaction in molecular dynamic simulations

A possible explanation for the inability of the SARS-CoV-2 ORF7a H47Y mutant to attenuate MHC-I levels on the surface of the cell is due to altered interactions with MHC-I. Molecular dynamic simulations have previously been used to describe the SARS-CoV-2 ORF7a-MHC interaction (17). We reasoned that similar simulations could be used to investigate for differences in binding to MHC-I between the WT and H47Y mutant ORF7a. We first performed protein-protein docking using ClusPro to model the interaction of WT and H47Y mutant ORF7a with MHC-I HLA-A2 (Fig. 5A). We then utilized CABS-flex 2.0 to determine changes in flexibility of both WT and mutant ORF7a bound to MHC-I (Fig. 5B). We observed two regions of altered structural flexibility, one of which was more flexible in ORF7a WT (between residues 19-23) and one that was more flexible in the ORF7a H47Y mutant (between residues 27-30). Interestingly, the region affected in the ORF7a H47Y mutant is within the predicted interface of ORF7a with MHC-I, which may alter the protein-protein interaction (Fig. 5A). This suggested that the H47Y mutation might compromise the ORF7a-MHC I interaction. We therefore performed molecular dynamic (MD) simulations using the WebGro server and determined the binding affinity (ΔG) trajectory of wildtype and H47Y mutant ORF7a in complex with MHC-I. Interestingly, we found that the binding affinity of ORF7a H47Y mutant with MHC-I appeared to be lower compared to that for the wildtype ORF7a over the course of MD simulation time (Fig. 5C). Together, these data show that the H47Y mutation of SARS-CoV-2 ORF7a may alter the architecture of the interaction between ORF7a and MHC I rendering it less efficient. In summary, the observed changes in protein dynamics may influence the functionality or stability of ORF7a like other single amino acid residue mutations previously identified (24).

**Fig. 5.**
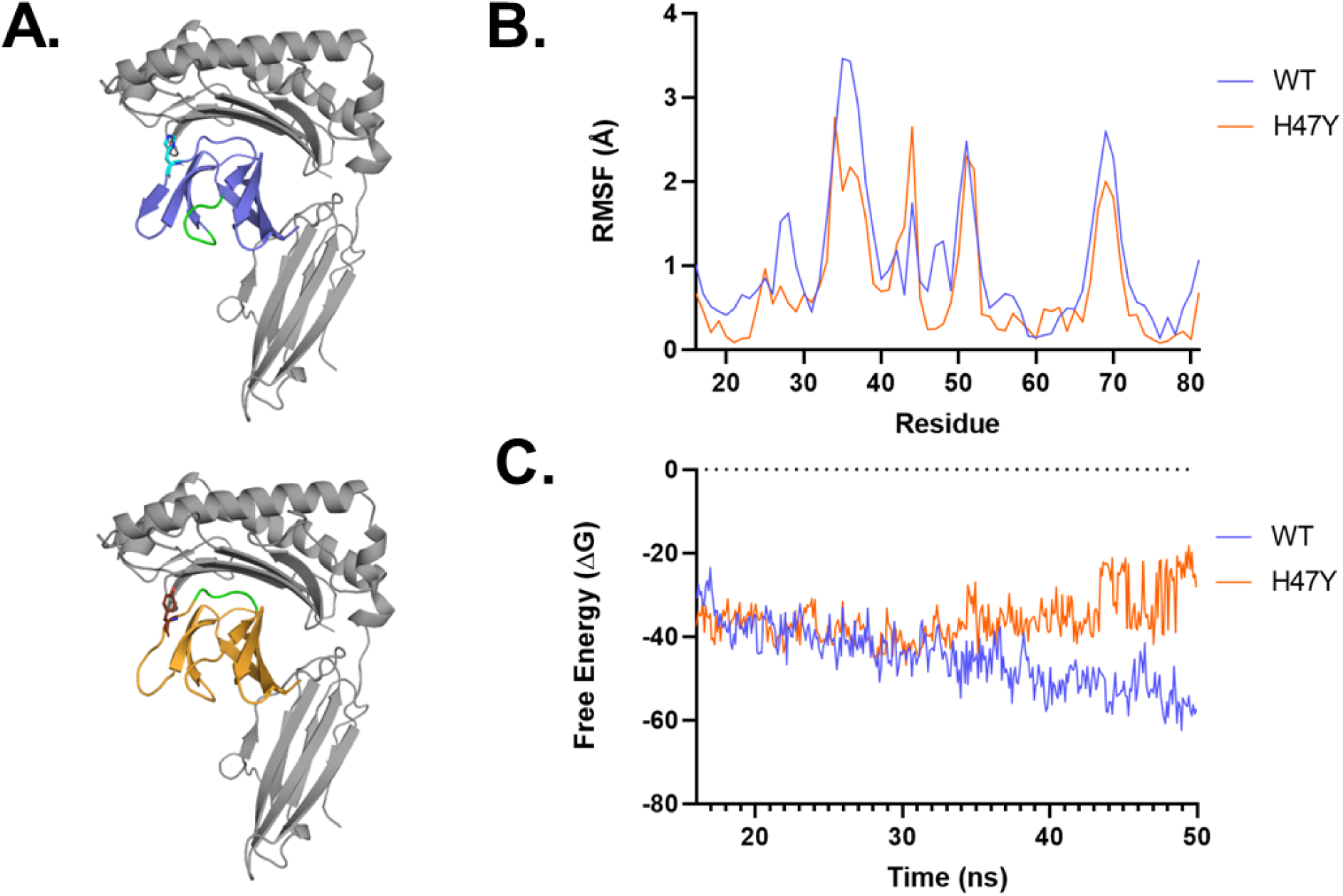
The H47Y mutation in SARS-CoV-2 ORF7a causes changes in protein structural dynamics. (**A**) Representative cartoon models from ClusPro docking simulations of wildtype (blue) or H47Y mutant (orange) SARS-CoV-2 ORF7a with MHC-I HLA-A2 (grey). Regions highlighted in green represent areas where increased flexibility was observed in the comparison between wildtype and mutant ORF7a. (**B**) Graphs depicting the root mean square fluctuation (RMSF, angstroms) calculated by CABS-flex 2.0 of the residue of wildtype and H47Y mutant SARS-CoV-2 ORF7a within the ORF7a-MHC-I complex. (**C**) Graphical representation of the average free energy trajectory values from 3 independent simulations of MHC-I with wildtype and H47Y mutant SARS-CoV-2 ORF7a.

## DISCUSSION

Herein, we focused on determining the effect of the ORF7a: H47Y mutation, prevalent among the SARS-CoV-2 Omicron subvariants BF.5 and BF.7, in various known functions of ORF7a, namely its anti-SERINC5 effect, its ability to antagonize the type I IFN response and its capacity to downregulate MHC-I cell surface levels.

We found that SARS-CoV-2 ORF7a carrying the H47Y mutation retains its ability to block the SERINC5-mediated restriction of SARS-CoV-2 infectivity. We have previously shown that deleting the SARS-CoV-2 ORF7a β-strands including the β-strand D, wherein the 47^th^ amino acid lies, has no effect on its anti-SERINC5 function which is governed by its TM domain (13). We speculate that with an unchanged TM domain, this ORF7a mutant is capable of interacting with SARS-CoV-2 S and SERINC5 similar to the SARS-CoV-2 ORF7a WT protein and hence able to block the antiviral effect of SERINC5.

Unlike SARS-CoV-2 ORF7a WT, our data showed that the H47Y mutation abolished the ability of SARS-CoV-2 ORF7a to suppress the type I IFN response. A single amino acid change in a protein may affect its own post-translational modifications as is the case for the R105G mutation in APOBEC3H (35, 36), so it is possible that the H47Y mutation may affect SARS-CoV-2 ORF7a ubiquitination as polyubiquitination of residue K119 has been reported to be critical for interfering with type I IFN response (15). More studies are needed to further elucidate the mechanism by which H47Y mutation interferes with the ability of SARS-CoV-2 ORF7a to counteract the type I IFN response. We also found that the H47Y substitution in SARS-CoV-2 ORF7a rendered it incapable of down-regulating surface MHC-I. Our in-silico analysis showed SARS-CoV-2 ORF7a H47Y substitution altered structural flexibility within the predicted interface of ORF7a with MHC-I likely altering the efficiency of protein-protein interaction. A recent study reported that Phe residue at position 59 (F59) in the E-F loop of SARS-CoV-2 ORF7a protein is a critical determinant for its ability to interact with MHC-I and retain it within the ER (18). It is interesting to note that H47Y site lies along a deep groove formed between the C-D and E-F loops of SARS-CoV-2 ORF7a protein and thus has the potential to alter the interactions of ORF7a with its cellular binding partners (12, 33). In fact, it is well established that naturally occurring mutations in viral proteins can have deleterious effects in their anti-MHC-I function. For example, polymorphisms within the HIV-1 accessory protein Negative Factor (Nef) modulate Nef-induced endocytosis of MHC-I from the cell surface (37). Moreover, a recent report identified that Omicron subvariants are able to downregulate MHC-I more efficiently from the surface of the cell than earlier isolates (38). In our studies, we did not examine the effect of these mutations during actual SARS-CoV-2 infection, because we found that the H47Y mutations affects primarily the ability of ORF7a to counteract the type I IFN response and the downregulation of MHC I from the cell surface; both events known to be targeted by multiple SARS-CoV-2 proteins (NSP1, NSP6, NSP13, M, N, ORF3a, ORF6, ORF7a, ORF7b) (15, 16, 39–41) and thus it is possible that the ORF7a H47Y effect may be masked by the function of the other viral proteins. In conclusion, our results show that the BF.7 and BF.5 associated H47Y mutations of SARS-CoV-2 ORF7a can modulate important function of this SARS-CoV-2 accessory protein, indicating that impairment of these functions may contribute to differences in viral pathogenesis.

## MATERIALS AND METHODS

### Data mining and reconstruction of phylogenetic tree

Surveillance data of 15,843,705 SARS-CoV-2 isolates were retrieved from the GISAID database on August 8, 2023 (https://www.gisaid.org) (42). We excluded the data of SARS-CoV-2 isolate that i) lacks PANGO lineage information; ii) was collected before July 31, 2023; iii) was isolated from non-human hosts; iv) was sampled from the original passage; and v) whose genomic sequence is not longer than 28,000 base pairs and contains ≥2% of unknown (N) nucleotides, resulting in the data of 1,943,768 SARS-CoV-2 isolates in the BA.5 lineage (EPI SET ID: EPI_SET_230817yu). At most ten genomic sequences of SARS-CoV-2 in each BA.5 subvariant (EPI SET ID: EPI_SET_230817cf) were randomly sampled and were subsequently aligned to the genomic sequence of Wuhan-Hu-1 SARS-CoV-2 isolate (NC_045512.2) using multiple pairwise alignment implemented in ViralMSA v1.1.24 (43). Gaps in the alignment were removed automatically using TrimAl v1.4.rev22 with-gappyout mode (44), and the flanking edges of the alignment at positions 1–388 and 29525–29713 were trimmed manually. A maximum likelihood-based phylogenetic tree was then reconstructed from the alignment using IQ-TREE v2.2.0 (45). The best-fit nucleotide substitution model was selected automatically using ModelFinder(46). Branch support was assessed using ultrafast bootstrap approximation (47) with 1,000 bootstrap replicates. We omitted a genomic sequence of Wuhan-Hu-1 from the reconstructed tree and rooted the tree using a genomic sequence of SARS-CoV-2 isolate whose tree distance was closest to the Wuhan-Hu-1 isolate.

The state of having or lacking ORF7a: H47Y substitution was assigned to terminal nodes of the reconstructed tree based on the mutation calling data from the GISAID database. The reconstruction of ancestral states was then performed using ace function of the ape R package v.5.7-1 (48) with equal-rate model. The ancestral node with a posterior probability of having the mutation at least 0.5 is considered having the mutation, whereas the node with the posterior probability less than 0.5 is considered lacking the mutation. The occurrence event of ORF7a: H47Y substitution was then determined from the state change from lacking mutation in the ancestral node to having mutation in the adjacent descendant node. The reconstructed tree was visualized using the ggtree R package v3.8.2 (49). All the phylogenetic analyses were aided by R v.4.3.1 (50).

### Multiple sequence alignment and generation of sequence logo plot

Genomic sequences of 78 sarbecoviruses, including those of SARS-CoV-1, SARS-CoV-1-related, SARS-CoV-2, and SARS-CoV-2-related viruses, were retrieved from the previous phylogenetic study (34). Each genomic sequence was aligned to each other using MAFFT v7.511 (51) with the G-INS-I mode and 1,000 maximum iterations. The coding nucleotide sequence of ORF7a was translated into the protein sequence using JalView v2.11.2.7 (52) according to the standard genetic code. Sequence logo plot for the ORF7a protein sequences was generated using WebLogo web service v3.7.12 (53) in default mode.

### Cell lines and maintenance

HEK 293T cells (ATCC, CRL-3216) and HEK 293T-hACE2 (BEI Resources, NIAID, NIH, NR-52511) were cultured in Dulbecco’s Modified Eagle Media (DMEM; Gibco) with 10% (vol/vol) fetal bovine serum (FBS; Sigma), and 100 mg/ml penicillin and streptomycin (Gibco) at 37 °C and 5.0% CO_2_. All cell lines were detached using 0.05% Trypsin-EDTA (Gibco) after washing once with phosphate buffer saline (PBS).

### Plasmids

The pBJ5-SERINC5-HA plasmid was obtained from Heinrich Gottlinger. The HIV-1 NL4-3ΔEnv-NanoLuc and pCMV SARS-CoV-2 SΔ19 were obtained from Paul Bieniasz. ISG56-Luc plasmid has been previously described and was a kind gift from Raymond Roos (54). Other plasmids used in this study include the pEGFP-N1 (Clonetech) and pRL-CMV vector (E226A) acquired from Promega. Cloning strategy for generating codon optimized SARS CoV-2 ORF7a in pCDNA-V5/His TOPO (Invitrogen) has been described previously (13). This codon optimized SARS CoV-2 ORF7a (WT) in pCDNA-V5/His TOPO plasmid was used as a template to generate SARS CoV-2 ORF7a: H47Y variant using the Phusion SDM kit (Thermo Fischer Scientific) and the following primers: H47Y_F; 5’-ACAAGTTCGCCTTGACGTG-3’ and H47Y_R; 5’-TGTCTGCAAGAGGGTAGAAGG-3’.

### Pseudovirus production

HEK 293T cells were seeded in a 6-well plate at a cell density of 0.5 × 10^6^ cells/well. Next day, cells were co-transfected with plasmids for HIV-1_NL4-3_ΔEnv-NanoLuc (2.5 μg), pCMV SARS-CoV-2 SΔ19 (0.73 μg), pBJ5-SERINC5-HA (0.5 μg) or pCDNA SARS-CoV-2 ORF7a-V5/His (WT/H47Y; 0.75 μg) or empty vector using Lipofectamine 3000 (Thermo Fisher Scientific) per manufacturer’s recommendation. 24 h after transfection, culture media was removed and replenished. Cells (see **Immunoblotting section**) and culture supernatants were harvested 48 h post transfection, processed as described previously (13) and used for infection experiments.

### Pseudovirus infectivity

For pseudovirus infectivity experiments, HEK 293T-hACE2 cells (2.5 x10^4^ cells/well) were seeded in a 96-well plate. Cells were infected the next day and lysed at 48 hpi. Luminescence was measured using Nano-Glo luciferase system (Promega) and a Biostack4 (BioTek) luminometer. Infectivity was determined by normalizing the luciferase signals to virus levels as determined by western blots probing for HIV-1 p24^CA^ on culture supernatants.

### Immunoblotting

Cells lysates were prepared as previously described (13, 55). The following antibodies were used for probing the blots: mouse anti-SARS CoV/SARS CoV-2 S (GeneTex), mouse anti-V5 (Thermo Fisher Scientific), rabbit anti-HA (Cell Signaling Technology, mouse anti-HIV-1 p24 (NIH/AIDS Reagent Program, ARP-4121), monoclonal anti-β-actin (Sigma-Aldrich), HRP-conjugated anti-rabbit IgG (Cell Signaling Technology) and HRP-conjugated anti-mouse IgG (EMD Millipore). Signals were detected using enhanced chemiluminescence detection kits Clarity and Clarity Max ECL (Bio-Rad) followed by quantitation of bands intensities using the ImageJ software (National Institutes of Health; https://imagej.nih.gov/ij/).

### IFN-I luciferase reporter assay

HEK 293T cells were seeded in a 12-well plate at a cell density of 0.25 × 10^6^ cells/well. Next day, cells were co-transfected with pISG56-Luc (100 ng), pRL-CMV (5 ng), pCDNA SARS-CoV-2 ORF7a-V5/His (WT/H47Y; 2 µg) or empty vector plasmids. At 16 h post transfection, cells were treated with 1,000 units/ml of human IFN-β (PBL assay science) or mock-treated (PBS). 8 h post-treatment, the cells were assayed for dual-luciferase activities using Dual-Glo Luciferase Assay System (Promega) and a Biostack4 (BioTek) luminometer. Cells from the duplicate wells of mock-treatment conditions were lysed and processed for immunoblotting for confirming SARS-CoV-2 ORF7a (WT/H47Y) expression (see **Immunoblotting section**).

### Cell surface MHC-I down-regulation assay

SARS-CoV-2 ORF7a-mediated downregulation of cell surface MHC-I levels were determined by flow cytometry as previously described with some modifications (17, 18). HEK 293T cells were seeded in a 12-well plate at a cell density of 0.25 × 10^6^ cells/well. Next day, cells were co-transfected with pEGFP-N1 (50 ng), pCDNA SARS-CoV-2 ORF7a-V5/His (WT/H47Y; 1 µg) or empty vector plasmids. 24 h post transfection, cells were detached from the plate and stained with anti-human HLA A, B, C-Alexa Flour 647 W6/32 (BioLegend) for 30 minutes at 4 °C. Cells were washed 2× with FACS buffer (PBS containing 2% FBS and 0.8 mM EDTA), fixed with 2% paraformaldehyde for 10 min at 4 °C, and acquired on BD LSRFortessa followed by analysis using FlowJo version 10.8.0. Cells from the duplicate wells were lysed and processed for immunoblotting for confirming SARS-CoV-2 ORF7a (WT/H47Y) expression (see **Immunoblotting section**).

### Protein-protein docking and molecular dynamic simulations

Protein docking was performed similarly to methods previously describe(17). Briefly, the crystal structures of SARS-CoV-2 ORF7a (PDBID: 6W37, residues 16-81) (56) and MHC-I HLA-A2 (PDBID: 1DUY) (57) were obtained from the protein data bank (PDB). The H47Y mutation on SARS-CoV-2 ORF7a was performed using the mutagenesis function in the PyMOL Molecular Graphics System, Version 2.5, Schrödinger, LLC. The structures of wildtype and H47Y mutant ORF7a with MHC-I were then submitted to ClusPro for docking simulations (58–60). To determine alterations in protein flexibility (Root mean square fluctuation, RMSF), the docked structures were input into CABS-flex 2.0 webserver using default conditions (61). Molecular dynamic simulations for ORF7a binding affinities was performed using the WebGro (62). 50 ns simulations with 1000 individual frames each were ran using a CHARMM 27 forcefield at 300 Kelvin, 1.0 bars of pressure, and at 0.15M NaCl. Change in free energy states were acquired for every 2 frames from 3 replicate simulations using gmx_MMPBSA (63, 64).

### Statistical analysis

Statistical analyses were performed using GraphPad Prism software version 9.5.0. Statistical tests used to determine significance are described in the figure legends. Comparisons with the p values less than 0.05 were considered to be significant.

## ACKNOWLEDGMENTS

We thank Amy Jacobs and Raymond Roos for generously providing reagents used in this study. The following reagent was obtained through the NIH HIV Reagent Program, Division of AIDS, NIAID, NIH: Monoclonal Anti-Human Immunodeficiency Virus Type 1 (HIV-1) p24 (AG3.0), APR-4121, contributed by Dr. Marie-Claire Gauduin. This work was supported by National Institutes of Health Grant R21 AI160950A1 to S.S.

## AUTHOR CONTRIBUTIONS

S.S. and K.S. conceived and directed the project. U.T. S.D. performed the experiments and analyzed data. A.P. performed phylogenetic analyses under supervision of J.I. U.T. and S.S. wrote and edited the manuscript.

## DECLERATION OF INTERESTS

Authors declare no competing interests.

## DATA AVAILABILITY

Surveillance datasets of SARS-CoV-2 isolates are available from the GISAID database (https://www.gisaid.org; EPI_SET_230817yu, and EPI_SET_230817cf). The supplemental table for each GISAID dataset is available in the GitHub repository (https://github.com/TheSatoLab/ORF7a_H47Y).

